# Phyllosphere fungal diversity generates pervasive non-additive effects on plant performance

**DOI:** 10.1101/2023.03.17.533234

**Authors:** Brianna K. Almeida, Elan H. Tran, Michelle E. Afkhami

## Abstract

- All plants naturally harbor diverse microbiomes that can dramatically impact their health and productivity. However, it remains unclear how microbiome diversity, especially in the phyllosphere, impacts intermicrobial interactions and consequent non-additive effects on plant productivity.
- Combining manipulative experiments, field collections, culturing, microbiome sequencing, and synthetic consortia, we experimentally tested for the first time how foliar fungal community diversity impacts plant productivity. We inoculated morning glories with 32 synthetic phyllosphere communities of either low or high diversity or with single fungal taxa, and measured effects on plant productivity and allocation.
- We found 1) non-additive effects were pervasive with 56% of microbial communities interacting synergistically or antagonistically to impact plant productivity, including some consortia capable of generating acute synergism (e.g., >1000% increase in productivity above the additive expectation), 2) interactions among ‘commensal’ fungi were responsible for this non-additivity in diverse communities, 3) synergistic interactions were ~4 times stronger than antagonistic effects, 4) fungal diversity affected the magnitude but not frequency or direction of non-additivity, and 5) diversity affected plant performance nonlinearly with highest performance in low microbial diversity treatments.
- These findings highlight the importance of interpreting plant-microbial interactions under a framework that incorporates intermicrobial interactions and non-additive outcomes to understand natural complexity.

## Introduction

Plants naturally harbor complex and diverse microbiomes containing many microbial taxa interacting with both their host plant and one another (Lundberg et al., 2012; Roman-Reyn et al., 2019; Trivedi et al., 2020). Plant-microbiome interactions shape a wide range of ecological processes (e.g. succession, community assembly, and speciation; Howard et al., 2020, Afkhami and Strauss 2016; Osborne et al., 2017) and contribute to many ecosystem services (e.g., carbon sequestration, nutrient cycling, and primary productivity (Averill et al., 2014; Gougoulias et al., 2014; Li et al., 2019; Harman et al., 2021). Much of our understanding of these interactions comes from decades of research and numerous manipulative experiments inoculating plants with individual microbial species of interest. These studies have repeatedly documented the importance of pathogens on plant health as well as how particular symbiotic microbes, such as *Epichloё* endophytes, rhizobia, and mycorrhizal fungi, can benefit plant productivity and fitness (Schardl 1996; Hoeksema et al., 2010; Thamer et al., 2011;) with cascading effects on the community dynamics of herbivores and other organisms (Hartley and Gange 2009; Arnold et al., 2014). For instance, systemic endophytic fungi living inside plants can improve host performance by priming plants against pathogens and reducing abiotic stresses, such as drought and nutrient limitation (Hubbard et al., 2014; Khare et al., 2018; Liu and Brettell 2019). While manipulative studies of single microbial symbionts have provided important insights into some of the microbial effects on plants, this approach typically ignores microbial diversity and thus does not consider the importance of intermicrobial interactions that undoubtedly are happening within the microbial communities for plant performance.

It is inherently challenging to test how microbial diversity and intermicrobial interactions within complex microbiomes impact plant performance. Some studies have embraced the natural complexity harbored within plant microbiomes by manipulating the presence of whole microbial communities (e.g. using ‘live’ soils as microbiome inoculum) (Morella et al., 2019; David et al 2020; Korenblum et al., 2020; Kiesewetter and Afkhami 2021). However, this method cannot easily determine which microbes and intermicrobial interactions underpin the observed changes in their host plants since the entire microbiome is changing at once. Recent synthesis has highlighted the importance of tripartite interactions as a bridge between single partner interactions and whole microbiome sequencing/inoculations as well as an avenue for studying the impact of microbial interactions on plant performance (Afkhami et al., 2020). Specifically, these tripartite studies have been leveraged to provide preliminary insights into how non-additivity can result from interactive effects of microbes within the microbiome. For instance, complementarity of the rewards or services that microbes provide to their host may lead to increased plant growth greater than the additive expectation (i.e. expected growth based on the benefits provided by each microbe alone) (Afkhami et al., 2014; Hori et al., 2021). For example, one study revealed a higher frequency of synergistic effects on *Panicum virgatum* (switchgrass) when inoculated with pairs of fungi that differed in functional traits that were likely to provide distinct and complementary rewards (Connor et al., 2017). While non-additive outcomes appear to be common based on tripartite studies (Larimer et al., 2010; Larimer et al., 2014, Ren et al., 2016), plant microbiomes are complex with many interactions between diverse taxa. Little is known about the frequency and importance of non-additivity for plant performance outcomes with more complex microbial communities and how diversity of the microbiome may impact non-additive responses. One pathway forward would be experiments manipulating diversity within synthetic microbial communities, which have served to experimentally link reductionist and community-level approaches to understanding microbial communities in the past (e.g. Niu et al., 2017, which used synthetic communities to demonstrate the impact of certain microbial taxa on bacterial community assembly in maize roots). Here, we embrace and manipulate phyllosphere microbial complexity using synthetic communities to determine how microbial diversity impacts non-additive outcomes of plant performance.

Diversity within microbial communities may affect the importance of intermicrobial interactions leading to non-additive effects on host plant productivity. In isolation microbes may act as either pathogens, mutualists, or commensals (harming, benefiting, or having no effect on plant health, respectively) toward their hosts, but their assumed role may change depending on the context in which the interaction takes place, such as in the context of high or low diversity microbial communities. The net effect of all cooperative and competitive interactions between microbes can lead to three possible outcomes for the plant (Afkhami et al., 2020). First, the host plant may perform better than the additive expectation from singly inoculated plants due to synergism of beneficial microbial effects. This outcome could result from complementary benefits provided by different microbes or cooperation among microbes. Second, the host plant may perform worse than the additive expectation from singly inoculated plants. This antagonism could result if members of the microbiome are in competition or conflict with one another. Third, the plant’s performance may match the additive expectation of singly inoculated plants, which could occur if microbes had no measurable impact on one another and the benefits or costs to the host. This outcome would also occur if synergistic and antagonistic effects among different groups of microbes within high diversity communities balance each other out or if neutral effects of many microbes in high diversity communities outweighs weak synergistic/antagonistic interactions.

In this study, we combined manipulative experiments, field collections, fungal culturing, microbiome sequencing, and synthetic microbial consortia to explore the importance of non-additivity and diversity in microbial effects on plant productivity. We inoculated plants with 16 high and 16 low diversity synthetic communities of foliar fungi and compared plant performance outcomes to additive expectations (based on single microbe inoculations) to determine whether changes in the phyllosphere microbiome’s diversity leads to synergistic, antagonistic or additive outcomes on plant performance. We determine 1) how frequently non-additivity occurs in microbial communities, 2) the consequences of microbial diversity for the strength and direction of non-additive effects, and 3) the impacts of foliar microbial diversity on host productivity. Our results highlight the pervasiveness of non-additive microbial effects on plant productivity and show that foliar fungal diversity influences the magnitude but not frequency of non-additivity, demonstrating that interactions within the host-associated microbiome can shape the interaction between plants and their microbiomes.

## Materials and Methods

### Study System and Field Collections

To determine how foliar fungal diversity impacts host productivity, we manipulated the leaf microbial community of *Ipomoea hederifolia,* a morning glory native to the Southeastern United States. *Ipomoea hederifolia* is readily found in disturbed upland habitat on the edges of the imperiled Pine Rocklands ecosystems. We collected leaves of *I. hederifolia* from 10 natural habitat fragments (16 ± 2.8 plants/site; total plants sampled=199) across Miami-Dade County to generate fungal isolates that form natural plant-fungal associations. Seeds for our experiment were collected from the Taylor R. Alexander Representative Species Assemblage of Natural Upland Communities of South Florida on the University of Miami campus (Coral Gables, Florida).

### Generating Culture Collection of Foliar Diversity

Following methods from Cook et al., (2013), a fungal culture library was created by placing the adaxial side of field collected leaves face down on potato dextrose agar plates, isolating individual emergent fungal isolates, and then propagating each isolate clonally. Antibiotics (ampicillin, streptomycin, or kanamycin) were used to prevent bacterial growth and to increase the diversity of fungal taxa acquired from collected leaves. Each fungal isolate was identified by extracting the fungal DNA and amplifying the ITS1-LR3 region using standard Extract-N-Amp protocols (David et al., 2016; Sigma Aldrich Corp., Saint Louis, Missouri). Samples were then sent to Eurofins (Louisville, KY, USA) for sanger sequencing, and the *BLASTn* algorithm was used to query NCBI databases for these sequences to identify the fungal isolates. To select which of these isolated and identified fungi would be used in the diversity treatments, we then sequenced the whole foliar fungal microbiomes from *I. hederfolia* leaves and selected the 20 unique fungal taxa from the culture collection that were most relatively abundant among the overall fungal communities (an indicator of dominance).Sequence data and plant performance data have been submitted to NCBI (PRJNA874722) and Zenodo (DOI 10.5281).

To profile the natural foliar fungal microbiome, we performed DNA extractions from a subset of our field collected leaves that were representative of the range of our collections (27 leaves) (Qiagen DNeasy Plant mini kit CatNo.69204) and used two-step dual indexing to prepare barcoded amplicon libraries of the ITS region (Gohl et al., 2016, Revillini et al., 2021) as well as a negative control library (using Ultrapure water in lieu of leaf tissue during extraction). Libraries were sequenced on the Illumina MiSeq platform (v3, paired end 300bp) at the University of Miami Center for Genome Technology. Sequences were demultiplexed (*bcl2fastq*), denoised and grouped into OTUs (operational taxonomic units) based on 99% similarity using the *QIIME2* (v3) pipeline and then rarified fungal sequences to 1000 reads at which rarefaction curves reached saturation. Taxa were identified with the UNITE database (ver. 01-12-2017), and the cultured fungal taxa were compared to fungi in the resulting community-wide taxa matrix to select which isolated fungi would be used in the experiment.

### Experimental Setup and Data Collection

To test the effects of fungal community diversity on plant productivity, we surface sterilized *I. hederifolia* seeds with 0.15% tebucanozole solution (as in Kucht et al., 2003) and then germinated the seeds in sterile petri plates with moist filter paper. Eight days after germination, seeds were aseptically planted in pots of sterile soil (262ml Heavyweight Deepots, Stuewe and Sons, Corvallis, Oregon) in the greenhouse at the University of Miami campus. Three weeks post-germination, leaves were abraded with sterilized sand and inoculated (Subedi et al., 2022) with either a single fungus from our culture collection (*i.e.,* monocultures), a ‘low’ microbial diversity treatment (*i.e.,* simple synthetic communities of 3 fungal taxa), a ‘high’ diversity treatment (*i.e.,* relatively more complex synthetic communities of 10 fungal taxa), or a sham inoculum of sterile water (*i.e.* control treatment with no microbial addition). We created inocula for each of the 20 fungal taxa (Christian et al., 2019), which were applied to 16 replicate monoculture plants per fungal isolate, and the sterile water control (sham ‘no microbe’ inoculum) was also applied to 16 plants. The two diversity treatments were generated by randomly selecting fungi from the pool of 20 isolated fungal taxa, resulting in 16 distinct low diversity synthetic communities and 16 distinct high diversity synthetic communities. Each of these 32 synthetic microbial communities were inoculated into three plants (Barrett et al., 2015). To create inocula, we suspended cultured fungal tissue of each fungus in 50 mL of sterile water, determined their concentrations using a cytometer, and diluted all fungal isolate solutions to equal concentrations. These solutions were used for monoculture inocula and to generate 15 ml of inoculum stock solutions of high and low diversity community treatments by combining the appropriate single isolate inocula in equal volumes (e.g. a low diversity inoculum combined 5 ml of each of three randomly chosen monoculture inocula). We also treated 16 plants with a leaf slurry inoculum made by aseptically homogenizing field-collected leaves containing the natural microbiome of *I. hederifolia* following Subedi et al., (2022) to ensure that the effects of synthetic fungal communities in our experiment were comparable to the microbial effects experienced by plants with natural foliar microbiome communities. Plants inoculated with synthetic communities performed similarly to plants inoculated with the natural microbiome via this slurry method with 91% of synthetic communities root mass and 81% shoot mass falling within one standard deviation of performance of leaf slurry (whole microbiome-treatment) plants. This suggests that the synthetic communities can provide a realistic model for microbial effects on plant performance.

The experiment was harvested five months after germination. Plant height (*i.e.,* length of the vine) as well as above and belowground biomass were measured. After root tissue was washed to remove soil, above and belowground biomass were dried separately at 60°C to a constant weight then measured (Mettler Toledo ME-T Analytical Balance, Columbus, OH). Total biomass was determined by taking the sum of above and belowground biomass.

### Statistical Analyses

Before performing statistical analyses, we assessed normality using Shapiro-Wilk tests and determined that log transformations were needed to improve normality for all performance data. To investigate if individual fungal taxa are mutualistic, commensal, or pathogenic, we compared the performance of monoculture inoculated plants to that of non-inoculated control plants with a MANOVA. To determine whether plant response to endophyte diversity is non-additive, we performed Monte Carlo simulations. A distribution of 9,999 plant trait values were created by randomly sampling values from the single inocula treatment plants that correspond to the fungal isolate combinations for each high and low diversity combination. Then 95% confidence intervals were determined from each distribution; if the actual mean plant performance from a high or low diversity treatment community fell outside of these intervals then the effect of fungal community combination on plant performance was determined to be non-additive (Crawford and Whitney 2010). We quantified how many times a foliar fungal community contributed to an antagonistic (≤ bottom 2.5% of the distribution) or synergistic effect (≥top 2.5% of the distribution) on any plant trait. We then investigated how likely it was for low or high diversity treatments to lead to antagonisms or synergisms on multiple plant traits using a chi-squared analysis with the frequency of antagonisms or synergisms as the response and diversity treatment as the predictor. To determine if the magnitude of positive or negative effects on plant performance varied between low and high diversity treatments, we first standardized each performance metric (standardized normal deviates) and calculated expected performance of plants grown with diversity treatments using the average performance values from plants grown with monoculture treatments. We performed an ANOVA with diversity treatment (i.e. low vs. high diversity synthetic communities) and direction of positive or negative effects compared to expected, as well as their interaction as categorical predictors and magnitude of effect (i.e., deviation in plant performance with diverse synthetic community from expected values based on additive effects from the monocultures) as the response variable.

To understand the relationship between foliar fungal diversity and plant growth, we analyzed the effects of fungal diversity on multivariate plant performance by performing a MANOVA with follow-up analyses on each fitness components using the same explanatory variables (after confirming significant overall microbial effects). The model included diversity treatments levels and synthetic community identity (nested within diversity treatments) as explanatory variables and plant growth metrics of root mass, shoot mass, and height as the response variables. We also conducted ANOVAs on plant investment (root-to-shoot ratio) or total biomass as response variables with diversity level and synthetic community identity nested within diversity level. Finally, we performed trend contrast analysis to evaluate the shape of the relationship between fungal diversity and plant growth. All statistical analyses were performed in R ver 3.6.0. (R Core Team, 2020).

## Results

### Fungal taxa functioned as commensals in isolation

We taxonomically identified the 20 foliar fungal isolates used in our experiment (Table 1), which included taxa that have been characterized in other systems as plant pathogens (e.g *Fusarium oxysporum, Colletotrichum gloeosporioides)* and saprotrophs (e.g. *Aspergillus sydowii)* as well as fungi characterized as having multiple lifestyles (e.g. *Sarocladium strictum* which has been characterized as a pathogen and saprotroph). When we compared the effect of single taxa inoculations to the control treatment, we found that all 20 fungi acted as commensal symbionts when inoculated alone (Fig. **1**), as they did not significantly change plant performance compared to the uninoculated control plants (e.g. Total Biomass F_20,139_=0.776, p=0.739; see Supplemental Table 1 for more details). Despite all these fungal isolates being commensal in pairwise interactions with the plant (*i.e.*, in monocultures), all fungal taxa were part of at least one synthetic community that had non-additive effects on plant performance. This result suggests that the presence of other fungi causes many commensals to shift their relationship with their plant host, resulting in changes to plant performance.

**Figure 1.**
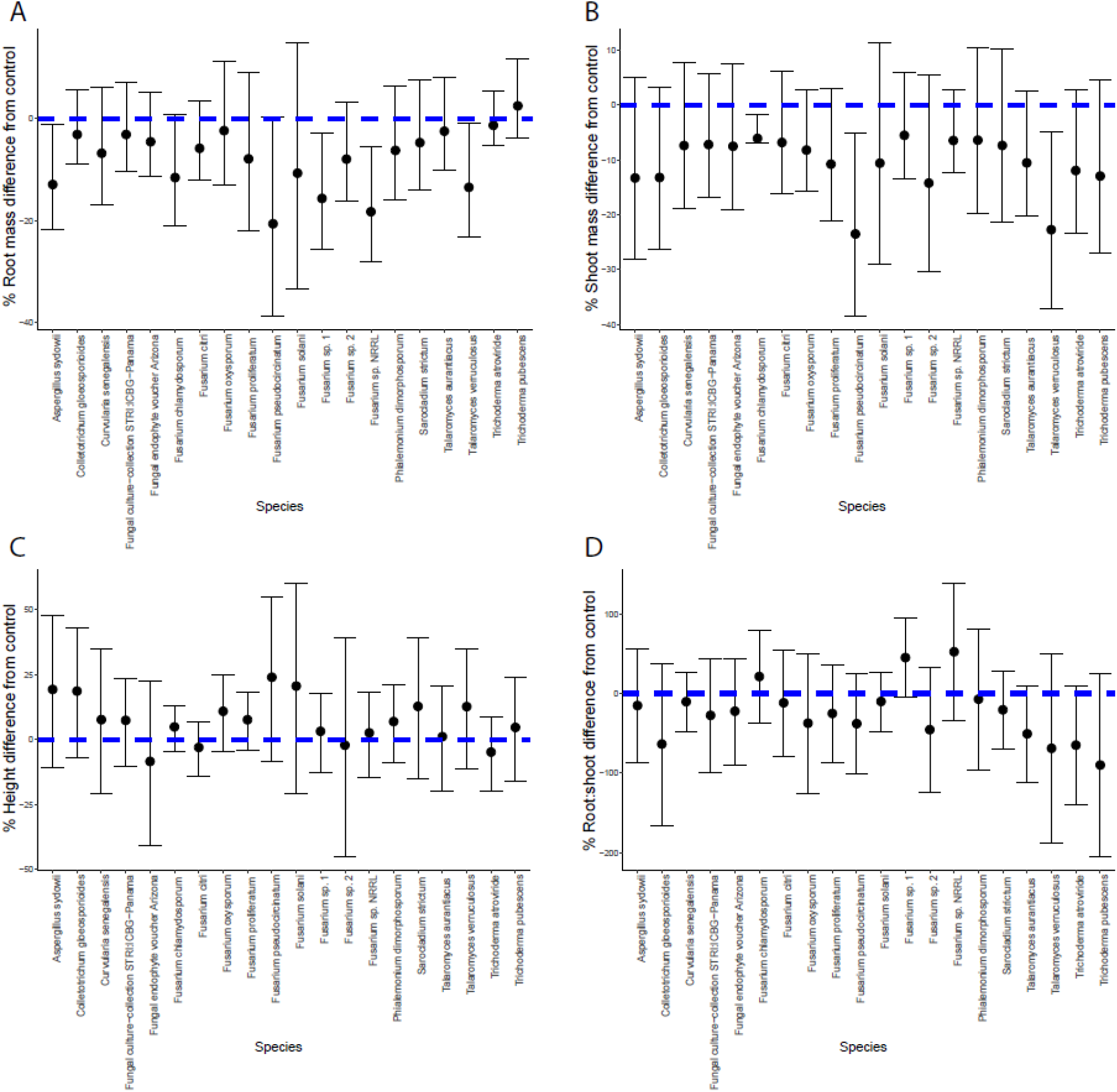
All fungal taxa used in the experiment acted as commensals when singly inoculated. This figure depicts the percent difference of the growth effects of each fungus compared to the control treatment (plants treated with sterile ‘sham’ inoculant) for (A) root biomass (B) shoot biomass, (C) height, and (D) investment in roots versus shoots (i.e. root-to-shoot ratio). Each point represents the mean percent difference for plants grown with one of the 20 fungal taxa, and error bars represent 95% CIs around the mean. Note that while the 95% Cis show a few taxa’s effects bounded away from 0 for biomass effects, the statistical results from ANOVA tests (available in Table S1) found that none of the fungi’s effects on plant performance were significantly different from the control plants. We use the results from the ANOVA over 95% CIs following best practices from previous diversity and productivity studies (Crawford and Whitney 2010) and because the ANOVA accounts for experiment-wide error rates.

**Table 1.**
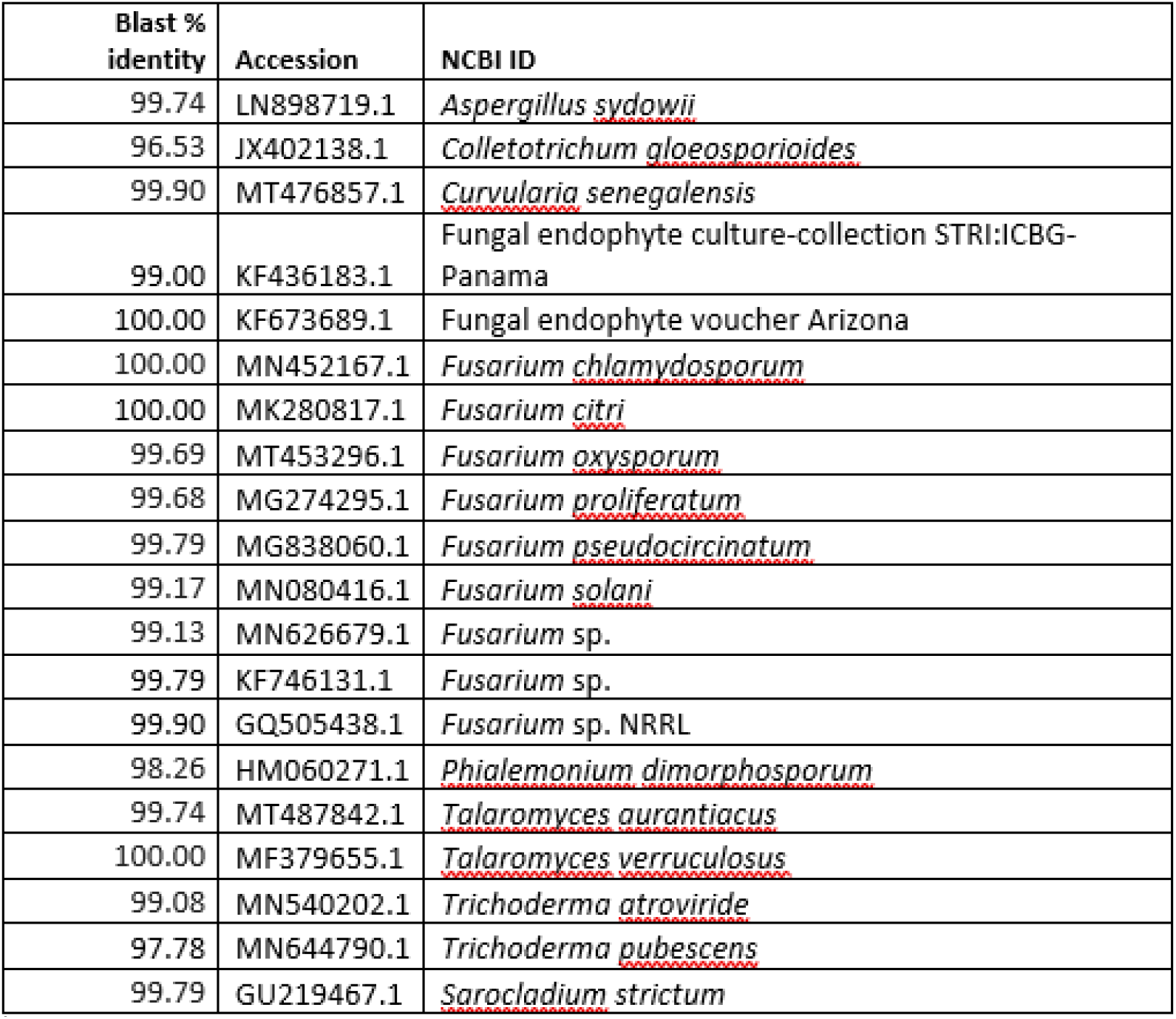
Information for the 20 fungi cultured, identified, and used in our synthetic communities. Note that Blast % identity refers to the extent to which sequences are aligned to the “NCBI ID” taxa sequences, and “Accession” refers to the NCBI accession numbers.

### Fungal communities composed of commensals have non-additive effects on plant performance

Monte Carlo analysis comparing performance of plants grown with synthetic fungal communities to the distribution of plant performances predicted based on single inocula treatments demonstrated that non-additive effects of fungal symbionts were ubiquitous. In fact, more than half of the synthetic communities (56% of 32 communities) had significant non-additive effects on at least one host plant performance metric and all four plant traits responded non-additively to at least one synthetic community (Fig. **2**). Further, both synergistic and antagonistic effects on plant performance (i.e. positive and negative non-additivity, respectively) were common, with 10 communities having synergistic effects and 9 having antagonistic effects at least once. We also noted several instances of “acute synergism”, including one case where plant productivity across all three growth metrics was over 1000% greater than the additive expectation based on the fungal effects on plant growth when in monoculture inoculations (Fig. **2**, community 3) and another fungal community that increased productivity by 350-600% across all productivity metrics (Fig. **2**, community 13).

**Figure 2.**
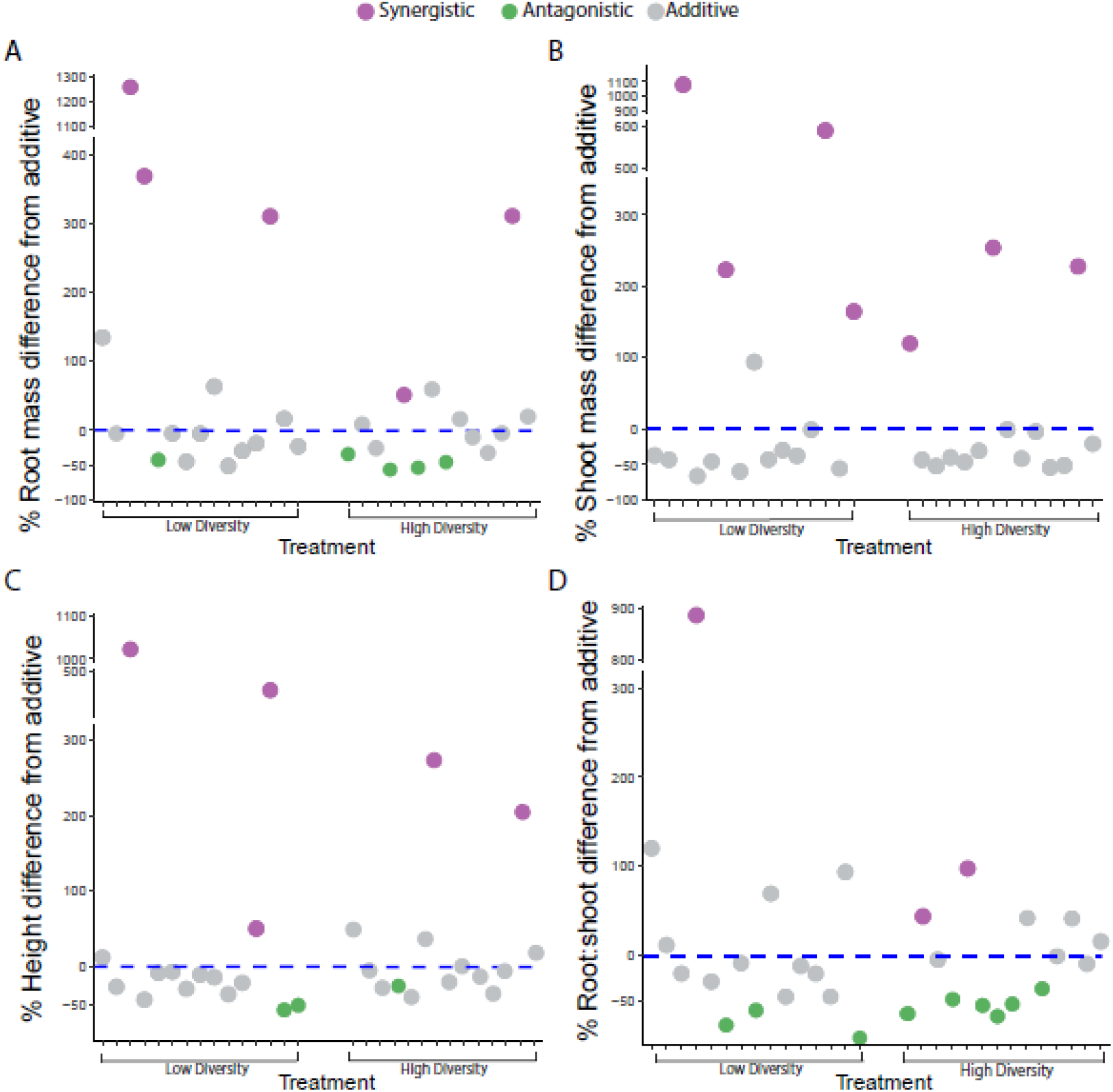
Non-additive effects of microbial communities on plant performance metrics of (A) root biomass, (B) shoot biomass, and (C) height as well as on (D) plant allocation between roots and shoots (root-to-shoot ratio). Each point represents the difference between a synthetic community’s effect on a given plant trait and the additive expected trait value calculated based on how the taxa that make up the synthetic community affected plants in the monoculture treatments (i.e. when singly inoculated). Purple points indicate synergistic community effects on plant productivity, green points indicate antagonistic community effects, and light gray points indicate additive community effects. The order of the low diversity and high diversity microbial communities is consistent across panels.

The probability of non-additive effects on plant growth did not depend on diversity of the microbial communities, as there was an equal number of high and low diversity communities causing non-additive outcomes on at least one plant performance metric (9 of 16 low diversity communities and 9 of 16 high diversity communities) (Fig. **2**). Further, when breaking non-additivity into synergistic and antagonistic effects on plant performance, we found that low and high diversity communities have an equal chance of non-additivity harming/benefiting plants (**χ^2^** = 0.29, p= 0.867, df=2). Specifically, there were 9 instances of significant synergistic effects on plant performance generated by 5 different high diversity communities and 11 instances generated by 6 different low diversity communities. Similarly, we found significant antagonistic effects in 11 instances across 7 high diversity communities and 6 instances with 5 low diversity communities (Fig. **2**). Interestingly, there were no negative non-additive effects on shoot mass in either low or high diversity treatment plants, but 7 instances of significant synergistic effects on shoot mass, suggesting beneficial non-additive outcomes from microbial interactions may be especially common in the part of the plant that this microbial community inhabits. We also note that while there was an equal frequency of synergistic and antagonistic non-additive effects generated by our synthetic consortia for most traits, synergistic effects were 4.24 (±16.1) times stronger on average than antagonistic effects (Fig. 2), suggesting beneficial intermicrobial interactions play an important role in microbiome-plant interactions.

We then investigated whether there was a difference in the strength of effects on plants of high and low diversity treatments using the deviation between the observed effects of synthetic fungal communities on plant performance and the expected performance (based on the average of the monoculture treatments). We found that the magnitude of positive and negative effects on host plants were significantly different when grown with high versus low diversity foliar fungal communities (F_1,112_=9.73, p=0.0023) (Fig. **3**). Low diversity treatments had a greater deviance from expected performance in both positive and negative directions, meaning that low diversity synthetic communities had stronger effects on plant growth compared to the higher diversity microbial communities. We also found that variance in this deviation from expected performance was greater for plants in the low diversity treatment than in the high diversity treatment (F_1,33_=4.87, p=0.029). Taken together this suggests that low diversity microbial communities may contain groups of interacting microbes that have substantive effects on plant performance, which are possibly diluted in the higher diversity treatment. This dilution may result from balancing of synergistic and antagonistic effects of different groups of microbes within the more diverse communities or neutral effects of many microbes outweighing any synergistic/antagonistic interactions.

**Figure 3.**
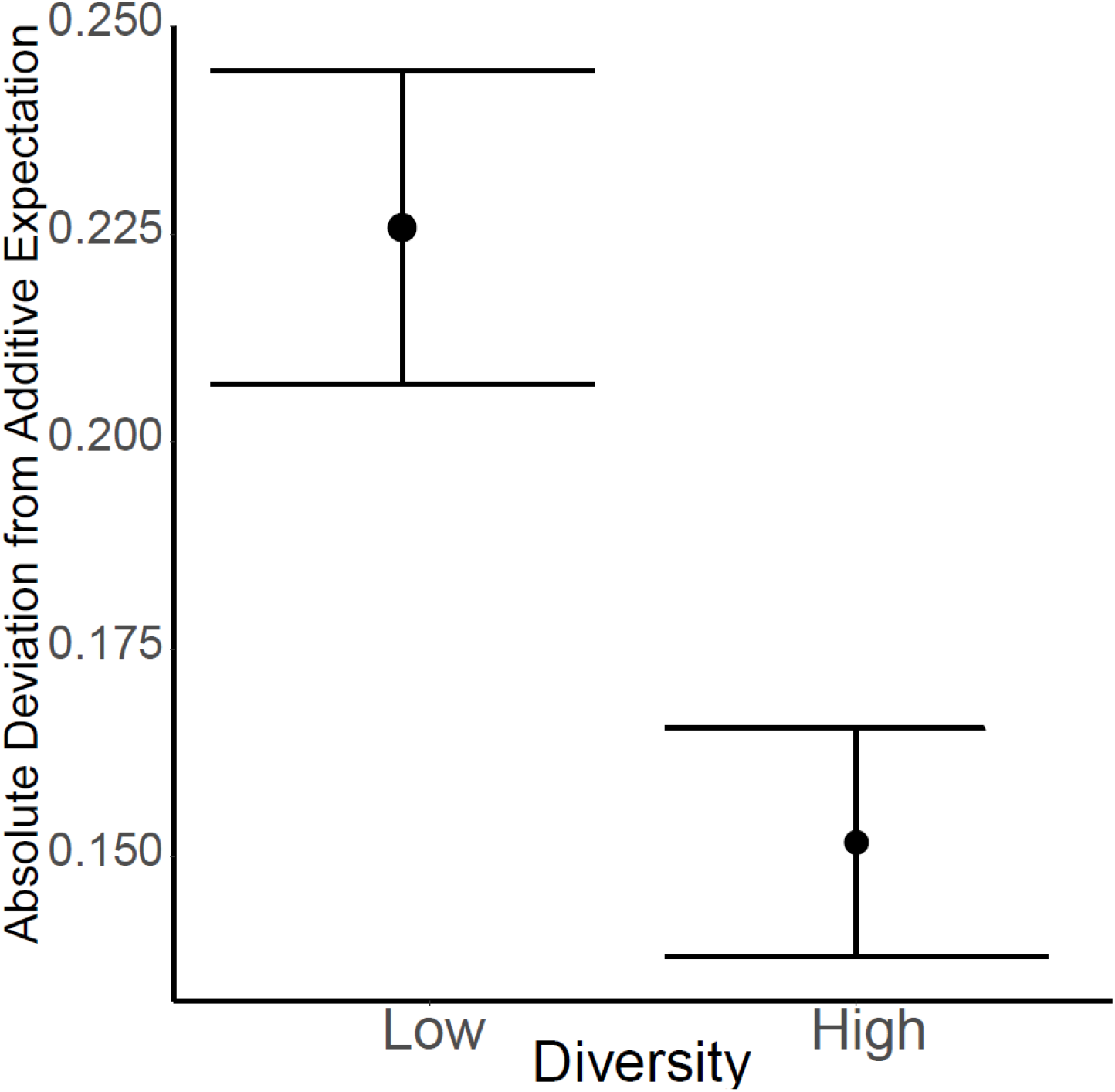
Increasing foliar fungal diversity decreased the magnitude of community effects among microbes on plant performance. We found that the deviation between plants inoculated with synthetic communities and the additive expectation was on average greater in the lower diversity treatment (Diversity effect: P=0.002). The deviation on the y-axis was determined by standardizing plant performance values (converting into standardized normal deviates to allow comparison of effect across traits that differ in scale) and calculating the difference between each synthetic community’s effect on plant performance and the additive expectation based on the monoculture performance outcomes. Points with bars indicate mean ± SE.

### Foliar fungal diversity impacts plant performance

The composition of the foliar fungal communities significantly affected overall plant performance (MANOVA: F_48,158_=1.28, p =0.029), and total biomass produced (F_48,158_=1.56, p=0.022). Diversity of the fungal communities was also important. For instance, root biomass showed a marginally significant increase with increasing foliar fungal diversity (F_3,158_=2.41, P=0.069; Fig. **4A**) and total biomass was significantly affected by diversity (F_3,158_=3.23, p=0.024; Fig. **4D**). Interestingly, the effect of fungal diversity on total biomass followed a quadratic relationship (trend contrast: t_204_=-2.11, p=0.036) with low diversity plants having the highest mass followed by high diversity and singly inoculated plants having the next highest, followed by sham-inoculated, control plants (Fig. **4D**). This demonstrates that plants can benefit from even very small phyllosphere communities compared to single species inocula, showing promise for the use of synthetic communities to promote plant growth even with the limitations of creating these communities.

**Figure 4.**
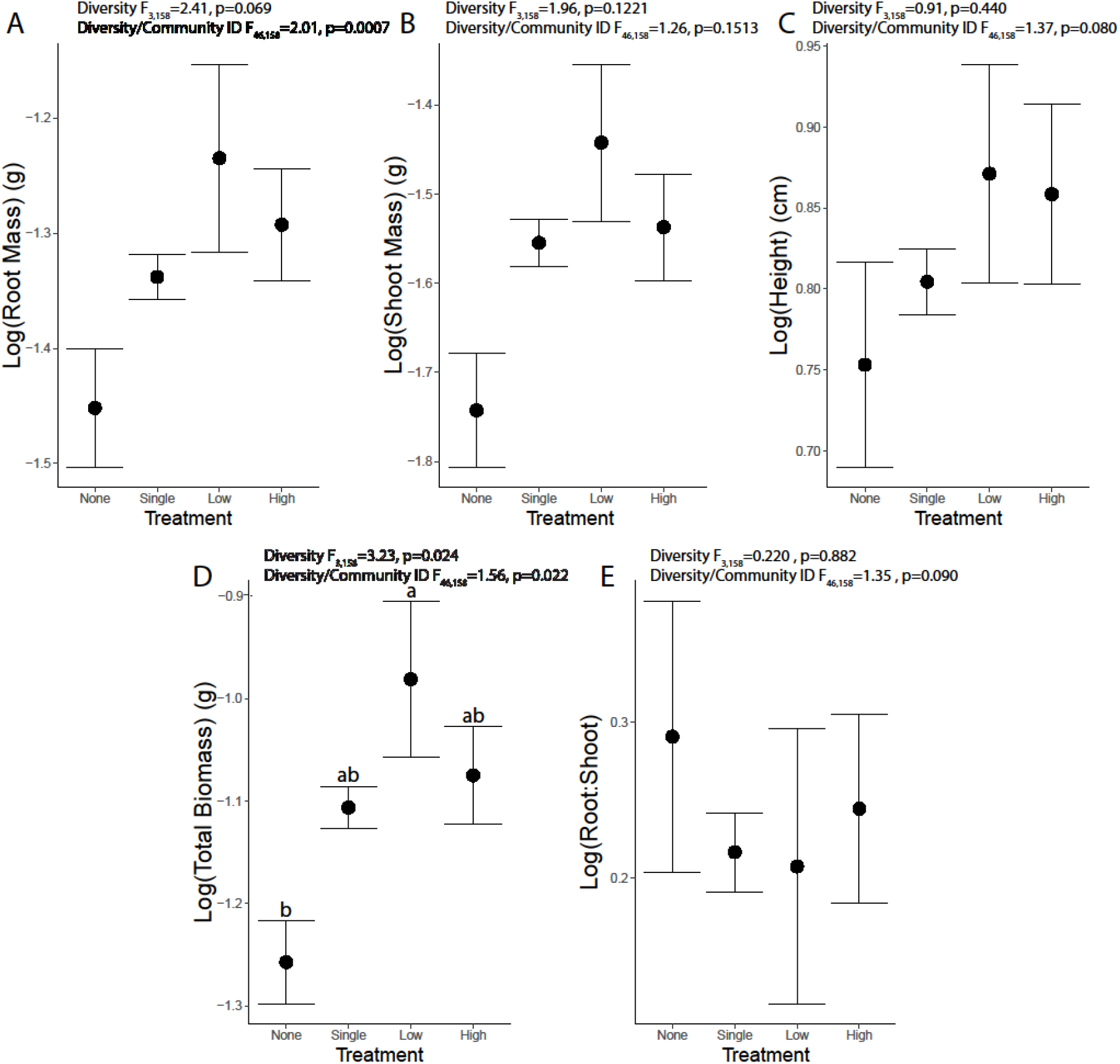
Effect of fungal diversity on host plant (A) root mass, (B) shoot mass, (C) height, (D) total biomass, and (E) the investment in roots versus shoots (i.e., root-to-shoot ratio). Points with bars indicate mean ± SE of log transformed data. Diversity/Community refers to community identity nested within diversity treatments.

## Discussion

To overcome the complexities of understanding whole microbial communities’ effects on plant performance, we used experimental synthetic communities to determine how increasing microbial diversity impacts the non-additive effects of microbes on plant performance. This study demonstrates five main results that contribute to our understanding of microbial diversity and non-additivity: 1) fungal isolates that act as commensals alone often have significant positive and negative effects on plant performance when in microbial consortia, 2) non-additivity is a common property of microbial communities with equal amounts of synergism and antagonism, 3) synergistic effects were often stronger than antagonistic effects on plant productivity, 4) the strength, but not the frequency, of non-additivity was affected by microbial community diversity, and 5) diversity’s effects on plant productivity followed a quadratic relationship, with the lower diversity communities having the greatest total biomass.

### Intermicrobial interactions among commensals can underpin pervasive non-additivity in plant-microbial interactions

All commensal microbes in our study participated in communities with non-additive effects on their host plant productivity, demonstrating interactions within microbiomes have consequences for plant performance. Our results also highlight that non-additivity is a pervasive feature of microbial communities as more than half of our randomly-generated synthetic communities had non-additive effects on at least one plant performance trait. Direct or indirect intermicrobial interactions could have led to this non-additivity through three (non-mutually exclusive) pathways. First, the presence of other community members changes the biotic context in which the interaction between a microbe and plant takes place, and this change in context can cause a microbe that is seemingly commensal in monoculture to confer benefits or act antagonistically when part of a natural microbial community. For example, a commensal microbe may have negative nontarget effects on host performance when engaging in competition with other microbes or conversely may become able to confer new or enhanced benefits to host plants when in the presence of other microbes that provide resources it needs. For instance, arbuscular mycorrhizal fungi can improve the ability of rhizobia to fix nitrogen by increasing the phosphorus uptake of the host plant (as N-fixation is a phosphorus-hungry process; Puschel et al., 2017). Second, the presence of other microbes in the community may alter the value of rewards or services that a microbe provides to its host plant. For example, many fluorescent foliar *Pseudomonas* spp. induce an immune response that can prevent infection by fungal pathogens (Van Wees et al., 2008), but the defensive effects of priming are only valuable to the host in the presence of pathogens. Third, multispecies interactions between microbes can provide novel functions for their host plant that lead to non-additive effects on plant growth. For example, co-inoculation of *Capsicum annuum* with *Acinetobacter* sp. and the putative plant pathogen *Phytophthora capsici* increased fresh weight of seedlings in an experiment. The researchers suggest this outcome resulted from changes in volatile organic compounds released by *Acinetobacter* in response to interactions with *P. capsici* that increased root length leading to greater fresh weight (Syed-Ab-Rahman et al., 2019). All three of these pathways are likely to play important roles in the non-additivity found in our study and in many other plant-microbiome interactions. Future work pinpointing when each mechanism is most important would be valuable for improving the predictability of microbial effects on plants and for applied goals requiring engineering of synthetic microbiomes in natural and human environments (De Souza et al., 2020; Sharm & Shukla 2020).

### Importance of microbial synergism and antagonistic effects on plant productivity

Using synthetic assemblies of microbes to understand the effects of microbial communities on plant performance (Grosskopf & Soyer 2014; Dolinsek et al., 2016), we show that not only is non-additivity ubiquitous but also that synergism and antagonism both occur often. While the frequency of these non-additive effects was the same regardless of diversity level, the strength of their effects on plant performance varied with microbial diversity level with stronger and more variable effects occurring in the low diversity treatment. Interestingly, synergistic effects were also ~4 times greater in magnitude on average compared to antagonistic effects, and we found a (non-significant) trend of positive effects of interactions within microbial communities being 10% greater than negative effects, on average, across all plant performance metrics and all synthetic communities. From the perspective of the host, these results indicate that not only is plant performance at least partially determined by non-additive intermicrobial interactions, but also that plants may experience stronger beneficial non-additive effects from microbial communities than negative non-additive effects. Importantly, our results documented multiple synthetic communities that had especially strong synergistic effects on plant productivity. For instance, one community increased all three plant productivity metrics - plant height, root biomass, and shoot biomass - by more than 1000% compared to the additive expectation and another microbial community increased productivity by 350-600% across all productivity metrics (see community 3 and community 13 in Fig. **2a**), emphasizing how important synergism among microbes can be for plant productivity. The community with the >1000% performance increase contained a fungal endophyte (previously detected in a tropical woody plant; vouchered accession KF436183.1), *Sarocladium strictum* (GU219467.1) and a species of *Fusarium* (KF746131.1; Table 1). *Sarocladium strictum* and *Fusarium* are often saprotrophs but also can act as plant pathogens suggesting that extreme synergistic effects can arise from interactions among taxa not typically considered plant mutualists (Rivera-Varas et al., 2007; Okungbowa &Shittu 2012). Given the strength of synergisms well beyond typical expectations for cooperative benefits, we hypothesize these synthetic communities with acute synergism are candidates for understanding novel functions driven by intermicrobial interactions. Future work functionally profiling synthetic communities like these could be valuable when considering microbiome engineering.

When compared with previous tripartite studies (between a plant and two microbes), non-additivity remains as important – if not more so – when plants interact with the more diverse communities used in our study. These synthetic communities also resulted in some plant performance outcomes that differ from tripartite studies. For instance, in contrast to the equal frequencies of synergism and antagonism we found, tripartite studies have often reported greater rates of synergism compared to antagonism when plants are inoculated with pairs of microbes (Larimer et al., 2014; Connor et al., 2017; Wezowicz et al., 2017) or in a few cases greater instances of antagonism on plant performance (Barrett et al., 2015; Ballhorn et al., 2016). However, the pattern of particularly strong synergistic effects in our study is in line with outcomes from tripartite studies where strength of synergistic effects compared to additive expectations has been as high as 238% and 180% (Larimer et al., 2014; Connor et al., 2017). Yet, the unprecedented strength of the acute synergism (e.g., >1000% compared to the additive expectation) generated by a few microbial consortia in our study far surpasses these previous highs from tripartite studies, prompting the need for future work manipulate diversity levels of synthetic consortia across different systems to determine how common these ‘acute synergism’ events are, what factors promote these outcomes, and what forms of direct or indirect inter-microbial interactions lead to them (e.g., working together to provide novel functions versus complementarity of rewards provided to the host by different microbes versus community enhancing a particularly important member of the microbiome, etc).

### Diversity-productivity relationships in phyllosphere plant-microbial interactions

A great deal of research has sought to generalize the relationship between community diversity and productivity, with much of the literature focusing on the relationship between plant diversity and plant productivity (Vermeer & Berendse 1983; Hector et al., 1999; Cadotte et al., 2009). These studies often find a positive relationship between plant community functional and phylogenetic diversity and plant community productivity. Leveraging a similar framework, researchers have become increasingly interested in how microbial community diversity could also shape host plant productivity. These studies have investigated belowground microbiomes, often finding a positive linear relationship between belowground fungal diversity and plant productivity (Van der Heijden et al., 1998; Van der Heijden et al., 2006; Vogelsang et al., 2006; Wagg et al., 2011; Koskella 2022). In contrast, our study – which focused on the phyllosphere microbiome diversity’s effects – documented a new quadratic relationship between foliar fungal diversity and total plant biomass. Additional diversity-productivity studies of the phyllosphere will be needed to assess if diversity of aboveground fungal communities consistently have different effects on host productivity than belowground communities. However, if common, systemic differences in effects of phyllosphere versus rhizosphere microbial diversity on host productivity could be related to the naturally lower diversity of phyllosphere microbiomes compared to rhizosphere microbiomes, which has been documented in studies of morning glories as well as other wild plants, crops, and invasive species (Dong et al., 2019; Zhao et al., 2019; Bao et al., 2020). There are several likely explanations for how intermicrobial interactions could underpin the quadratic relationship we found. For instance, the non-linear relationship could result from the neutral effects of many microbes in high diversity communities diluting the effects of smaller modules of synergistically or antagonistically interacting microbes on plant productivity. Alternatively, some synergistic or antagonistic interactions among fungi in the low diversity treatment could be disrupted by interactions with other taxa in high diversity communities (e.g., microbially secreted allelopathic chemicals harming a taxon integral to a synergistic interaction; Afkhami et al., 2020; Brown et al., 2020). These interactions within the fungal community may be contributing to the non-additive effects we see in this study leading to plant performance in which the highest diversity treatment plants are comparable to singly inoculated plants.

### Conclusions and perspectives on new directions

Overall, this research highlights how interactions within host-associated microbiomes can strongly and non-additively affect the interaction between plants and microbes as well as the promise of synthetic microbial consortia to understand the complex plant-microbial interactions that drive host health and productivity. Through this work, we have also identified several areas that would build on and complement our findings. In particular, the frequency of non-additivity and the complexity of the microbiome necessitates future work that evaluates the mechanisms that underpin non-additive effects on host productivity and the inter-microbial interactions involved. Differentiating among indirect and direct pathways is intrinsically challenging, especially since multiple mechanisms can simultaneously impact the net effect of the microbiome on the host. To address this challenge, we suggest future work that uses meta-transcriptomic approaches aimed at characterizing changes in microbial functional gene expression between microbes inoculated singly and in more diverse communities as well as co-expression networks analysis to identify shifts in expression that change in tandem between the microbial community members and plant hosts (Palakurty et al., 2018). High-throughput single-cell transcriptomics would further increase the ability to disentangle microbial interactions by allowing evaluation of inter-microbial co-expression of functional genes (Libault et al., 2010; Ma et al., 2019; Mauger et al., 2022) and could be especially useful for understanding how each microbe and their interactions contribute to the acute synergism events we found for several consortia in this study. Understanding expression of these functional genes could also provide insight into how species’ niche overlaps interact with increasing community diversity to impact plant performance across many functional dimensions simultaneously (e.g., greater functional diversity underlying the effects of greater species diversity on plant productivity; Connor et al., 2017; Afkhami et al 2020). In conclusion, our study shows promise for the use of synthetic microbial consortia to understand the complex plant-microbial interactions that underpinning plant health and microbial community structure and highlights new avenues for future investigations of microbial diversity and intermicrobial interactions.

## Acknowledgments

Thanks to K. Djahed for his assistance with experiment maintenance, and K. Crawford for advice on diversity-productivity study methods. Thanks to D. Hernandez, K. Kiesewetter, A. Igwe, A. Rawstern, B. Whitlock, and C. Searcy as well as the Wilson, Silveira and Mueller Lab for feedback on this manuscript. Thanks to the Environmentally Endangered Lands Program of Miami Dade County for allowing sample collections and the National Science Foundation for support to M. Afkhami (NSF DEB-1922521 and NSF DEB-2030060) and to B. Almeida (NSF Graduate Research Fellowship Program).

## Author Contributions

B. Almeida and M. Afkhami designed the research; B. Almeida and E. Tran established the experiment and collected the data. B. Almeida analyzed the data with guidance from M. Afkhami. B. Almeida, E. Tran and M. Afkhami wrote and edited the manuscript.

## Data Availability Statement

Data will be made available through NCBI Sequence Read Archive and Zenodo upon acceptance of the manuscript.

---

**Table S1.** ANOVA table for analysis showing that there was not a significant difference between the plant performance of control plants and the single strain inoculated plants across any of the performance metrics, indicating that all 20 fungal taxa were effectively commensals.

| **Response** | **Df** | **SS** | **Mean SS** | **F** | **P** |
| --- | --- | --- | --- | --- | --- |
| Root Mass | 20,139 | 1.15 | 0.057 | 1.08 | 0.377 |
| Shoot Mass | 20,139 | 1.36 | 0.068 | 0.617 | 0.895 |
| Height | 20,139 | 0.676 | 0.033 | 0.516 | 0.956 |
| Total Biomass | 20,139 | 0.943 | 0.047 | 0.776 | 0.739 |

